# CPGL: Prediction of compound-protein interaction by integrating graph attention network with long short-term memory neural network

**DOI:** 10.1101/2022.04.19.488691

**Authors:** Minghua Zhao, Min Yuan, Yaning Yang, Steven X Xu

## Abstract

Recent advancements of artificial intelligence based on deep learning algorithms have made it possible to computationally predict compound-protein interaction (CPI) without conducting laboratory experiments. In this manuscript, we integrated a graph attention network (GAT) for compounds and a long short-term memory neural network (LSTM) for proteins, used end-to-end representation learning for both compounds and proteins, and proposed a deep learning algorithm, CPGL (CPI with GAT and LSTM) to optimize the feature extraction from compounds and proteins and to improve the model robustness and generalizability. CPGL demonstrated an excellent predictive performance and outperforms recently reported deep learning models. Based on 3 public CPI datasets, C.elegans, Human and BindingDB, CPGL represented 1 - 5% improvement compared to existing deep-learning models. Our method also achieves excellent results on datasets with imbalanced positive and negative proportions constructed based on the above two datasets. More importantly, using 2 label reversal datasets, GPCR and Kinase, CPGL showed superior performance compared to other existing deep learning models. The AUC were substantially improved by 15% to 50% on the Kinase dataset, indicative of the robustness and generalizability of CPGL.

## Introduction

Binding affinity is a parameter describing interaction of a drug with a target protein, and can be used to predict in vivo pharmacological effects (efficacy or/and safety) of drug candidates ^12,21,23^. In drug discovery, potential drug candidates are often screened for binding affinity or compound-protein interaction (CPI). However, in vitro techniques measuring binding affinity in laboratories are usually expensive and time-consuming ^32^.

Recent advancements of artificial intelligence based on deep learning algorithms have made it possible to computationally predict CPI without conducting laboratory experiments^31^. Convolutional neural network (CNN) based models have been developed to extract features of compounds and proteins ^18,28,29^. To improve the learning of the representation of drugs, graph neural networks (GNNs) and graph CNNs^14^ (GCNs) have been proposed to extract structural features from the 2-D structure of compounds ^3,27,34^. In addition, recurrent neural networks (RNN) were also used for feature extraction from compounds and proteins ^1,6,11,40^.

Long-short term memory (LSTM) is a recurrent neural network designed to process sequence information^8,10^. Compared with CNN, LSTM has been shown to better capture the temporal information of input data and the relevant information with the large gap between the relevant input^38^. That is, LSTM is capable of learning the relationship between words that are far apart in the sequence. This unique utility of LSTM may allow better extraction of spatial features of protein structure. Conversely, CNN based approaches may only process adjacent amino acids in the sequence for feature extraction.

In addition, the 2D structure of small molecule compounds (i.e., atoms and chemical bonds) can be naturally viewed as a graph structure, where each node corresponds to an atom, and each edge corresponds to a chemical bond between 2 atoms. CNN designed for image data usually does not work well for molecule graph data which does not possess a Euclidean structure^41^. GNN, particularly graph attention network (GAT) is designed to process the information from graph structures and can learn the strength of the connection (i.e., chemical bonds) between different nodes (i.e., atoms) at the same time. Therefore, GAT can markedly improve feature extraction for small molecule compounds.

In this manuscript, we proposed CPGL (Compound-protein interaction prediction with graph neural network and long short-term memory neural network) to optimize the feature extraction from compounds and proteins by integrating the GAT for compound structures with LSTM for protein representation. We demonstrated that, via coupling with end-to-end learning, CPGL could improve the prediction of CPI on multiple datasets. More importantly, CPGL markedly improved the generalizability of model prediction, and could provide much greater prediction accuracy on label-reversal datasets compared to other recently published deep-learning algorithms (e.g., as CPI-GNN^34^, GCN^14^, GraphDTA^27^, and TransformerCPI^3^, etc).

## Methods

The proposed CPGL method is mainly composed of two deep learning models: the GAT model for molecular graph and the bi-directional LSTM for protein sequence (see Fig.1 for an illustration of the CPGL method). We will first review these two models (GAT and LSTM) in the two subsequent subsections and describe the CPGL model in the compound-protein interaction prediction subsection. Training method of the proposed model is briefly described in the modeling training and validation subsection.

**Figure 1:**
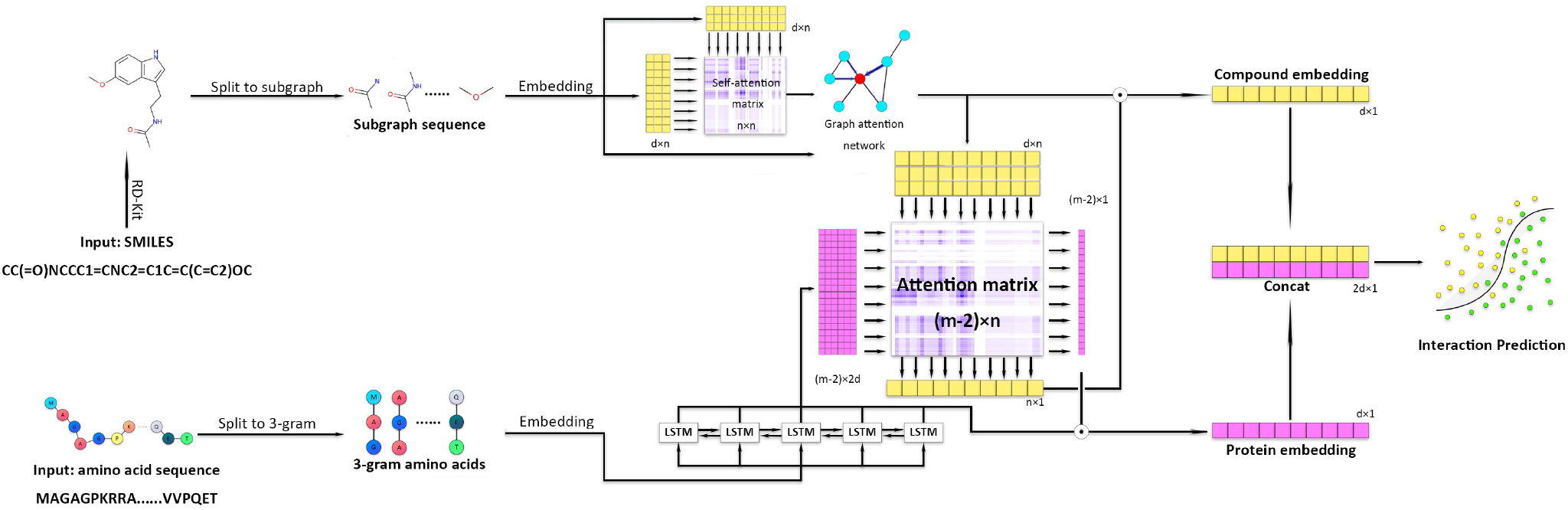
An overview of the CPGL. The proposed method is based on end-to-end representation learning, which uses raw inputs of compounds and proteins (i.e. SMILES and amino acid sequences) instead of features such as molecular fingerprints and protein domains extracted from databases. The input molecular graphs are obtained from SMILES by preprocessing with RDKit while protein sequences are split based on n-gram amino acids (d is the vector dimension, n is the length of the atom and m is the length of the amino acid sequence). Compound and protein vectors, which are obtained using a GAT and a LSTM, respectively, are concatenated through a two-sided attention mechanism, and used as the input for a classifier to predict the compound-protein interaction.

### Graph attention network for molecular graph

We denote a molecular graph by 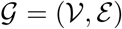, where 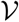 is the set of vertices representing the atoms in the molecular and 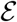 is the set of edges representing the chemical bonds in the molecular. Let *v_i_* be the *i*-th atom in 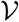 and *e_ij_* be the chemical bond between the *i*-th atom and the *j*-th atom. We use *r*-radius subgraph^4^ to reinforce representation learning. More precisely, we denote 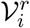 to be the subgraph centered at the *i*-th atom with the atoms linked with atom *i* by at most *r* chemical bonds and the chemical bonds linking them, and 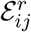 to be the subgraph centered at chemical bond with the atoms connected with *e_ij_* by at most *r*-1 chemical bonds and the chemical bonds linking them.

By using the toolkit RDKit^16^, we convert SMILES to *r*-radius subgraphs sequence. After random initialization, we represent 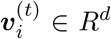 as the 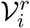 embedding at iteration *t*. Denote ***A*** as the adjacency matrix of the whole graph. We next introduce the transition function for atoms and chemical bonds.

Assume that there are *n* atoms in a molecule. For *i, j*=1,…,*n*, we calculate the graph attention weights as follows,

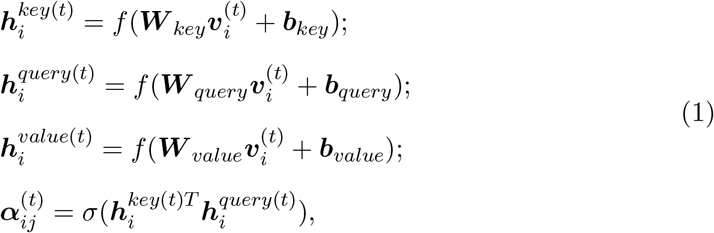

where *f, σ* are non-linear activation functions such as ReLU (^17^) or tanh function, ***W**_key_, **W**_query_, **W**_value_* ∈ *R^d×d^* are the weight matrices, and ***b**_key_, **b**_query_, **b**_value_* ∈ ***R**^d^* are the bias vectors. The attention weights 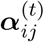 are used to choose the direction of information enrichment. Inspired by electron cloud distribution of molecules, the density difference of electron cloud distribution between atoms is described through the attention matrix.

Then we update the whole subgraph embedding vector sequence as follows,

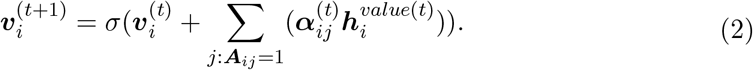

Thus, with the transition function (1) and (2) of graph attention network, we get compound vector sequence ***v***_1_, ***v***_2_,…,***v**_n_* as the output of the molecule network.

### Bi-directional LSTM for protein sequence

The LSTM model is a deep neural network designed to capture long-term dependence structure in natural language context. Since amino acids that are close in space may be far away in terms of text sequence representation, we choose to capture the spatial structure by applying the LSTM to process amino acid sequence information. Since the direction of an amino acid sequence is irrelevant in compound-protein interaction, we use the bi-directional LSTM model in our study.

Before applying the bi-directional LSTM to proteins, we first use word2vec algorithm to embed the amino acid sequence into real-valued vectors (Kimothi at al., 2016; Kobeissy et al., 2015; Mazzaferro., 2017;Yang et al., 2018). We then split the amino acid sequences into overlapping pieces or “words” of *k*-gram amino acids^5^. In this study, we set the length of the word to be *k* = 3. For example, we split *MAAVRM…LDLK* into “*MAA*”,“*AAV*”,…, “*DLK*”. Given an *k*-gram amino acid sequence, we randomly embed them into a vector sequence ***x***_1_,***x***_2_,…,***x***_*m*−*k*+1_ as the input of the bi-directional LSTM, where *m* is the length of amino acid sequence and ***x**_i_* ∈ *R^d^* is the embedding of the *i*-th word obtained by the pretraining approach word2vec ^25,26^. The output of the bi-directional LSTM will be represented by ***s***_1_,***s***_2_,…,***s***_*m*−*k*+1_, where ***s**_i_* ∈ *R*^2*d*^.

### Compound-protein interaction prediction

Given compound vector sequence ***V*** = (***v***_1_,***v***_2_,…,***v**_n_*) and protein vector sequence ***S*** = (***s***_1_,***s***_2_,…,***s***_*m*−*k*+1_), we introduce the attention matrix between compound vector sequence and protein vector sequence as follows,

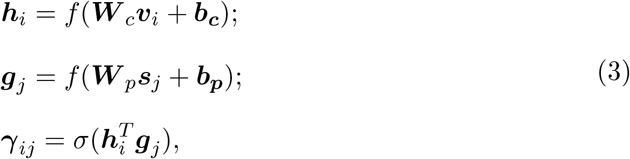

where ***W**_c_* ∈ *R^d^×^d^*, ***W**_p_* ∈ *R*^*d*×2*d*^ are the weight matrices and ***b**_c_, **b**_p_* ∈ *R^d^* are the bias vectors. Define weighted sum of ***V*** and ***S*** as follows,

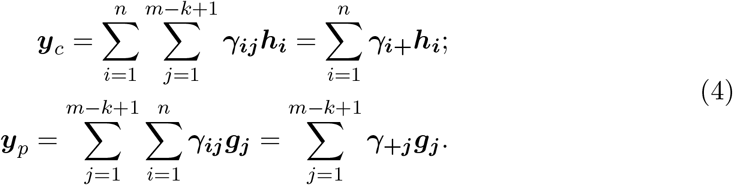

The two-sided attention mechanism can give different weights to different parts of the compound and protein, which allows us to pay more attention to the parts that have a greater impact in the binding. This allows us to model the interaction between the compound and the protein instead of obtaining a simple summation. Note that the attention matrix also allows us to analyze and find the key parts of compound and protein.

To predict the interaction, we concatenate ***y**_c_* and ***y**_p_* and obtain an full-connected network as follows,

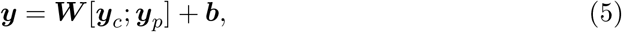

where ***W*** ∈ *R*^2×2*d*^ is the weight matrix, ***b*** ∈ *R*^2^ is the bias vector. Finally, we apply a softmax function on the final output vector ***y*** = [*y*_0_,*y*_1_] to compute the probability of class of interaction (1 for interaction, 0 for no interaction) as follows,

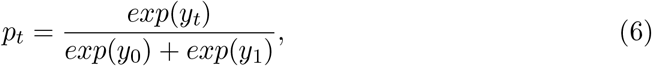

where *t* ∈{0, 1} is the binary label and *p_t_* is the probability output from softmax function for interaction class *t*.

### Modeling Training and Validation

To evaluate the performance of the algorithm and increase its stability, we used crossvalidation. The data set was randomly shuffled and partitioned into 6 parts, the first part of which was used as the test set, and the other part was used as the training set for 5-fold cross-validation, where one fold is used as the validation set and the rest is used as the training set. The final model was the average of these 5 models. All the settings and hyper-parameters of GPCL are summarized in Table 1.

**Table 1:** Hyper-parameters of CPGL

| Hyper-parameter | Value | order |
| --- | --- | --- |
| Radius($r$) | | 0,1,2 |
| Vector dimension($d$) | | 16,32,64,128 |
| Number of GAT layer |  | 3 |
| Number of output layer |  | 3 |
| Regularization |  | 1e-5,1e-6,1e-7 |
| Batch size |  | 1 |
| n-folds validation |  | 5 |

With known compound-protein pairs and the interaction in the training dataset, we use the *L*2-penalized cross-entropy as loss function:

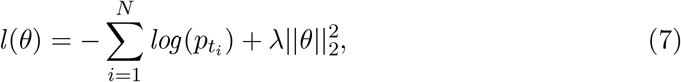

where *θ* is the set of all parameters in our model, *N* is the number of compound-protein pairs in the training dataset, *t_i_* is the interaction of the *i*-th pair, and *λ* is the hyper-parameter of *L*2 penalty. We used backpropagation algorithm to train our model.

GPCL was implemented with pytorch 1.7.1 and we used the LookAhead ^39^ optimizer combined with RAdam ^20^, which does not suffer from the divergence problems of Adam optimizer without the learning rate warmup^3^.

## Materials

### Public datasets

We used three public datasets, for human, C.elegans^19^ and BindingDB^6^ to compare the performance of our model with other machine learning and deep learningbased approaches. Positive samples with CPI of human dataset and C.elegans dataset were obtained from two manually curated databases: DrugBank 4.1^36^ and Matador^9^. Validated negative samples of compound-protein pairs are also included in the two dataset^19^. In total, 3,369 positive interactions among 1,052 unique compounds and 852 unique proteins are present in the human dataset, while 4,000 positive interactions among 1,434 unique compounds and 2,504 unique proteins are included in the C.elegans dataset. Since most negative protein-compound pairs that have no CPI have not been experimentally validated or reported in the literature, we used the validated negative samples screened by a computer program in Liu. et.al 2015^19^. The rationale behind this method is that proteins that are dissimilar to the known target of a given compound are unlikely to be the target of the compound. BindingDB dataset contains 39,747 positive examples and 31,218 negative examples with 49,745 unique compounds and 812 unique proteins from a public database^7^.

Since the proportion of positive and negative samples in real data is often unbalanced, we also synthesized unbalanced datasets from the human and C.elegans datasets to evaluate the robustness of the model. The ratio of positive to negative samples was set to be 1,3 and 5, respectively, and the training, validating and testing sets were randomly partitioned^34^. The training and testing data sets were carefully designed so that they have no common ligands or protein in CPI pairs. Therefore, BindingDB dataset can assess models’ generalization ability to unknown ligands and proteins.

### Label reversal datasets

Label reversal datasets were proposed to evaluate the robustness and generalizability of deep learning models^3^, where a ligand in the training set appears only in one class of interaction (either positive or negative interaction pairs), whereas, in the test set, the same ligand appears only in the opposite class of interaction. Two label reversal datasets,GPCR and Kinase, created by Chen et al., 2020 were used in this analysis to further evaluate model performance. Table 2 summarizes all datasets we use.

**Table 2:** Summary of datasets

|  | Proteins | Compounds | Interactions | Poisitive | Negative |
| --- | --- | --- | --- | --- | --- |
| Human | 852 | 1,052 | 6,728 | 3,369 | 3,359 |
| C.elegans | 2,504 | 1,434 | 7,786 | 4,000 | 3,786 |
| BindingDB | 812 | 49,745 | 70,965 | 39,747 | 31,218 |
| GPCR | 356 | 5,359 | 15,343 | 7,989 | 7,354 |
| Kinase | 229 | 1,644 | 111,237 | 23,190 | 88,047 |

## Results and discussion

### Performance on the balanced public datasets

We compared CPGL with existing machine learning models (i.e., K nearest neighbors (KNN), random forest (RF), L2-logistic (L2), and support vector machines (SVM)), and recently published deep learning-based models i.e., GCN^14^, CPI-GNN^34^, DrugVQA^40^, TransformerCPI^3^, GraphDTA^27^ (by tailoring the last layer to binary classification task) on human and C.elegans (Table 3 and Table 4) datasets. We followed the same training and evaluating strategies as CPIĺCGNN and repeated with ten different random seeds to evaluate CPGL. Area Under Receiver Operating Characteristic Curve (AUC), precision and recall of each model are compared.

**Table 3:** Comparison results of the proposed model and baselines on human dataset

| Method | AUC | Precision | Recall |
| --- | --- | --- | --- |
| KNN | 0.860 | 0.927 | 0.798 |
| RF | 0.940 | 0.897 | 0.861 |
| L2 | 0.911 | 0.913 | 0.867 |
| SVM | 0.910 | **0.966** | **0.969** |
| GraphDTA | 0.960±0.005 | 0.882±0.040 | 0.912±0.040 |
| GCN | 0.956±0.004 | 0.862±0.006 | 0.928±0.010 |
| CPI-GNN | 0.970 | 0.918 | 0.923 |
| DrugVQA | 0.964±0.005 | 0.897±0.004 | 0.948±0.003 |
| TransformerCPI | 0.973±0.002 | 0.916±0.006 | 0.925±0.006 |
| CPGL | **0.979±0.001** | 0.915±0.010 | 0.957±0.008 |

**Table 4:** Comparison results of the proposed model and baselines on C.elegans dataset

| Method | AUC | Precision | Recall |
| --- | --- | --- | --- |
| KNN | 0.858 | 0.801 | 0.827 |
| RF | 0.902 | 0.821 | 0.844 |
| L2 | 0.892 | 0.890 | 0.877 |
| SVM | 0.894 | 0.785 | 0.818 |
| GraphDTA | 0.974±0.004 | 0.927±0.015 | 0.912±0.023 |
| GCN | 0.975±0.004 | 0.921±0.008 | 0.927±0.006 |
| CPI-GNN | 0.978 | 0.938 | 0.929 |
| TransformerCPI | 0.988±0.002 | 0.952±0.006 | 0.953±0.005 |
| CPGL | **0.990±0.001** | **0.956±0.003** | **0.957±0.005** |

For the C.elegans dataset, CPGL demonstrated an excellent performance, and outperformed all the studied existing machine learning and deep learning models in terms of all the 3 evaluation metrics (AUC, precision, and recall). The AUC of CPGL is 0.99, while the precision and recall of CPGL are both around 0.96. This represents approximately 0.6 - 0.8% improvement compared to the most recently published deep learning algorithm, TransformerCPI. CPGL demonstrates 2 - 5% improvement compared to other recently developed deep learning-based approaches.

In terms of the human dataset, CPGL improved the AUC by 1%, compared to TransformerCPI. Greater degree of improvements is observed compared to other deep learning algorithms. In general, for both human and C.elegans datasets, the deep learning models had better performance compared to conventional machine learning based approaches. It should be noted that models based on 3D structural information of protein are out of scope for this manuscript as such information is not available for these two datasets. In general, for both human and C.elegans datasets, the deep learning models had better performance compared to conventional machine learning based approaches.

Since the traditional methods such as KNN,L2,SVM and RF are generally not comparable to various deep learning methods, we only compare our model with other deep learning methods for BindingDb dataset. Area Under Precision Recall Curve (PRC) and AUC of each model are shown in Table 5 and CPGL outperformed all the deep learning models in terms of AUC and PRC. The AUC of CPGL is 0.985 and the PRC is 0.983. This represents approximately 3.6% improvement compared with TransfomerCPI. We conclude that CPGL is still stable even in the face of unknown proteins and compounds. It should be noted that models based on 3D structural information of protein are out of scope for this manuscript as such information is not available for these three datasets.

**Table 5:** Comparison results of the proposed model and baselines on BindingDB dataset

| Method | AUC | PRC |
| --- | --- | --- |
| GraphDTA | 0.929 | 0.917 |
| GCN | 0.927 | 0.913 |
| CPI-GNN | 0.603 | 0.543 |
| TransformerCPI | 0.951 | 0.949 |
| CPGL | **0.985** | **0.983** |

### Performance on unbalanced public datasets

Furthermore, we evaluated CPGL using unbalanced datasets (i.e., different ratios of positive to negative samples) which was synthesized based on public datasets, human and C.elegans, and compared CPGL with KNN, RF, L2, SVM, CPI-GNN and TransformerCPI using the unbalanced datasets. Fig. 2 showes the AUC scores on the human and C.elegans unbalanced datasets. It is apparent that CPGL achieved the best perdictive performance on all the synthesized data regardless of degree of unbalance. Compared to TransformerCPI and CPI-GNN, CPGL improved the AUC by 1.8 - 3.3% and 0.6 - 2.9%, respectively. The performance of traditional machine learning models such as SVM, L2, RF, and kNN was markedly lower compared to CPGL.

**Figure 2:**
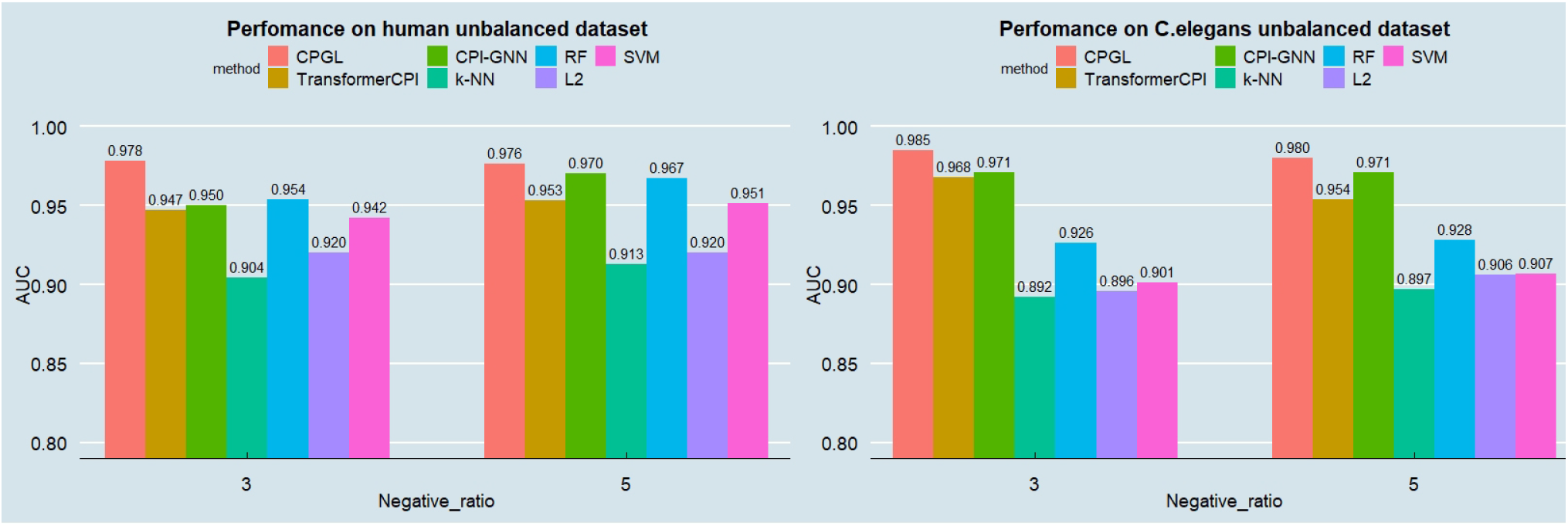
The AUC scores of various methods on the human and C.elegans unbalanced datasets

### Performance on label reversal datasets

We also used two label reversal datasets^3^, GPCR and Kinase, to compare our proposed CPGL with CPI-GNN, GraphDTA, GCN and TrasformerCPI. As shown in Fig. 3, On the GPCR set, CPGL have achieved similar AUC (0.856 vs. 0.863) but slightly better PRC (0.861 vs. 0.859) compared to TransformerCPI and outperformed other method both in AUC and PRC. At the same time, our method has made substantial improvement in kinase dataset. Compared with TransformerCPI, the AUC and PRC of CPGL are increased by about 17% and 22%. Meanwhile, the AUC values of other methods are less than 0.5, which indicates that these methods have failed on the kinase dataset. It is also known that most ligands in Kinase dataset possess almost ten times more noninteraction pairs than interaction pairs ^3^, while most ligands in GPCR dataset has relatively balanced positive and negative samples. This showed that our method is not subject to the hidden ligand bias and possesses capability of learning interactions between proteins and compounds. Overall, the results on label reversal dataset show that CPGL is more robust under the label reversal datasets and unbalanced datasets.

**Figure 3:**
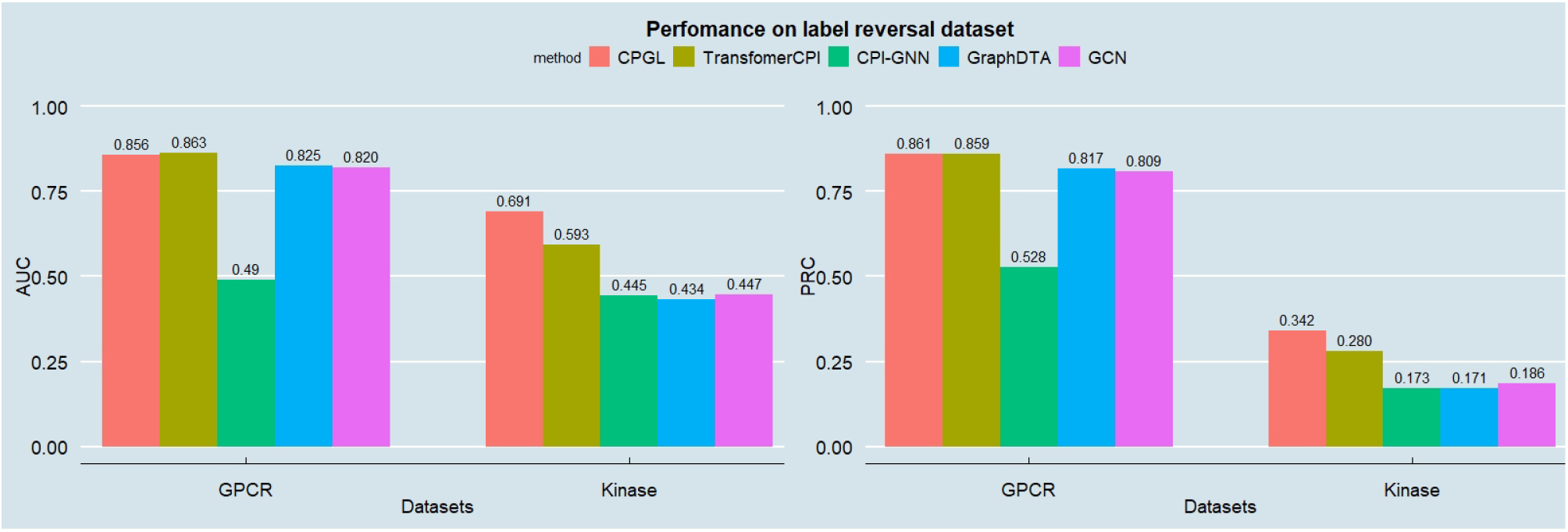
Results of CPGL, TransformerCPI, CPI-GNN, GraphDTA, GCN and CPGL without graph attention, CPGL without two-sided attention on GPCR dataset and Kinase dataset

### Model ablation study

In order to assess if the two attention mechanisms we used in the model are necessary, we evaluated two reduced models by removing the graph attention or the two-sided attention component from CPGL respectively using the label reversal experiment. As shown in Table 6, although the result of CPGL without graph attention on GPCR dataset are similar to those of CPGL with attention, it has a decrease of 9.0% and 23.1% on AUC and PRC, respectively, on kinase dataset. Meanwhile, the AUC and PRC of CPGL without two-sided attention decreased by 2.3% and 2.6% respectively on GPCR dataset and 5.9% and 21.3% respectively on GPCR dataset. These results show that both attention components in CPGL are beneficial.

**Table 6:** Model ablation study

| Method | GPCR |  | Kinase |  |
| --- | --- | --- | --- | --- |
|  | AUC | PRC | AUC | PRC |
| CPGL | 0.856 | 0.861 | 0.691 | 0.342 |
| CPGL w.o. two-sided-attn | 0.836 | 0.839 | 0.650 | 0.269 |
| CPGL w.o. graph-attn | 0.855 | 0.864 | 0.629 | 0.263 |

## Conclusion

We developed a deep-learning based algorithm, CPGL to optimize the feature extraction from compounds and proteins by integrating the GAT for compound structures with LSTM for protein representation. CPGL not only improved the prediction on multiple regular public datasets, but also substantially outperformed other reported deep-learning algorithms on unbalanced datasets and label-reversal datasets, indicative of the robustness and generalizability of CPGL.

## Data and code availability

The data and source codes of CPGL are available on the GitHub repository at https://github.com/RobinDoyle/CPGL.

